# Mathematical bridge between epidemiological and molecular data on cancer and beyond

**DOI:** 10.1101/2022.09.07.507053

**Authors:** Saumitra Chakravarty, Khandker Aftarul Islam, Shah Ishmam Mohtashim

**Affiliations:** Department of Pathology, Bangabandhu Sheikh Mujib Medical University, Shahbagh, Dhaka, 1000, Bangladesh; Department of Computer Science and Engineering, Bangladesh University of Engineering and Technology, Dhaka, 1000, Bangladesh; Department of Chemistry, Purdue University, West Lafayette, 47907, Indiana, USA

**Keywords:** Mathematical Modeling, Carcinogenesis, Epidemiology

## Abstract

**Background:** At least six different mathematical models of cancer and their countless variations and combinations have been published to date in the scientific literature that reasonably explain epidemiological prediction of multi-step carcinogenesis. Each one deals with a particular set of problems at a given organizational level ranging from populations to genes. Any of the models adopted in those articles so far do not account for both epidemiological and molecular levels of carcinogenesis.

**Methods:** We have developed a mathematically rigorous system to derive those equations satisfying the basic assumptions of both epidemiology and molecular biology without incorporating arbitrary numerical coefficients or constants devoid of any causal explanation just to fit the empirical data. The dataset we have used encompasses 21 major categories of cancer, 124 selected populations, 108 cancer registries, 5 continents, and 14,067,894 individual cases.

**Results:** We generalized all the epidemiological and molecular data using our derived equations through linear and non-linear regression and found all the necessary coefficients to explain the data. We also tested our equations against non-neoplastic conditions satisfying equivalent mathematical assumptions.

**Conclusion:** The aim of this treatise is not only to provide some novel insight into the mathematical modeling of malignant transformation but also to revive the classical tools we already have at our disposal to pave the way towards novel insight into integrated approaches in cancer research.

## 1. Introduction

We have built our derivation of the system of equations on the basis of the classical ‘log-log linear’ models proposed by Armitage [1], elaborated by Burch and used by Knudson [2] [3] to formulate his famous ‘two-hit hypothesis’. [4] [5] [6] After meticulously examining the possible interpretations of that model for each of the quantities within the equations, we have come up with a mathematically rigorous system to derive those equations that satisfy the basic assumptions of both epidemiology [7] and molecular biology without having to incorporate arbitrary numerical coefficients or constants devoid of causal explanation just to fit the empirical data. The dataset we have used encompasses 21 major categories of cancer, 124 selected populations, 108 cancer registries, 5 continents, and 14,067,894 individual cases. Our system of equations is therefore entirely algebraic. Then we developed variations of the log-log linear model that addresses the conditions posed by selective growth advantage/disadvantage and variable mutation/epimutation rates at different age groups. The developed model is based on some assumptions which are listed as follows:

### 1.1. Assumptions

Unless otherwise specified, following assumptions are held throughout the article:

1. Each type of cancer is the outcome of a series of discrete irreversible rate-limiting events, number of such events is denoted as ‘r’.
2. The individual rate-limiting events have very low probability that allows most people to live up to typical average human lifespan without ever developing any cancer.
3. Sequence of such rate-limiting events for a given cancer is unique, although the components of the sequence and the order may vary for different cancers in different individuals.
4. All of the rate-limiting events for a given cancer have to occur in an individual for the cancer to manifest.
5. Once the exact sequence of rate-limiting events for a given cancer has orchestrated in an individual, development of that cancer is inevitable.
6. There is a negligible time lag between all the events being executed appropriately to give rise to a cancer and the clinical or symptomatic manifestations of the cancer.
7. Rate of a given rate-limiting event is constant and time-invariant in each of the individuals in a susceptible population but different such events may have different rates.

For the rest of the article, in order to refer to any of the assumptions, we would simply mention the assumption number in the preceding text.

### 1.2. Models

Following the given assumptions, three distinct mathematical models of cancer are proposed:

1. Linear log-log model
2. Convex upwards log-log model
3. Concave upwards log-log model

The derivations of these models can be found in the Appendix.

#### 1.2.1. Linear log-log model

The linear log-log model is given by

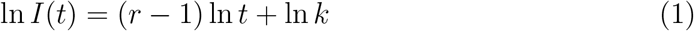

where *I*(*t*) is the age-specific incidence rate of cancer at age *t, r* is the number of rate-limiting events and *k* is the probability of *r* rate-limiting events to occur in a specific order within the given unit of time *t*. This form has the advantage of being linear on log-log plot where the tangent (*r* − 1) gives the direct measure of the number of driver mutations *r* required for cancer development, i.e., *y* = *mx* + *c*, where *y* = ln *I*(*t*), *x* = ln *t, m* = *r* − 1 and *c* = ln *k*.

#### 1.2.2. Convex upwards model

The convex upwards model is given by

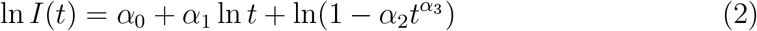

where *α*_0_, *α*_1_, *α*_2_ and *α*_3_ contains the following terms for the two different assumptions

##### Convex upwards model from heterogeneity assumption

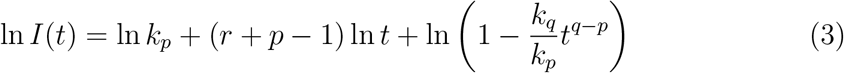

##### Convex upwards models from age-related effect assumption

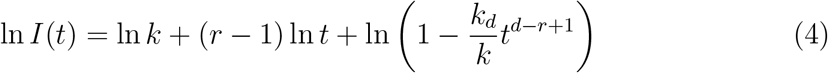

#### 1.2.3. Concave upwards model

The concave upwards model is given by

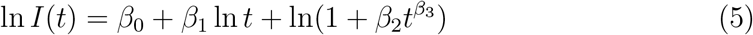

where *β*_0_, *β*_1_, *β*_2_ and*β*_3_ contains the following terms for the two different assumptions.

##### Concave upwards model from heterogeneity assumption

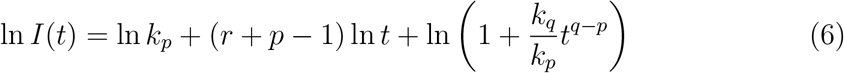

##### Concave upwards models from age-related effect assumption

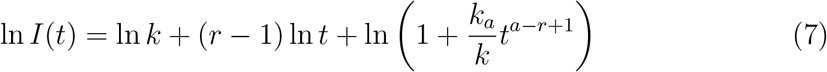

## 2. Methods

### 2.1. Data

We have tested our model on 21 categories of cancers listed in the GLOBOCAN database [17], of 124 selected populations from 108 cancer registries published in CI5 (Cancer Incidence in Five Continents) of the age groups: 0-14, 15-39, 40-44, 45-49, 50-54, 55-59, 60-64, 65-69. The registry contained the data for 14,067,894 people. The findings were correlated with the molecular data given in the latest editions of the World Health Organization (WHO) reference series on tumors published by the International Agency for Research on Cancer (IARC).

### 2.2. Statistical Analysis

Analysis was performed using R [18] and graphs were made using ggplot of R. For the linear log-log model a linear regression algorithm was used on the data collected from GLOBOCAN repository. For the non-linear (convex upwards and concave upwards model) a simple hybrid algorithm was used: Linear regression algorithm for the distinct linear part of the data, and an additional algebraic calculation for the additional term which explains the upwards curve in the later stages of the age groups. The related files have been uploaded to a GitHub Repository. [19]

## 3. Results

### 3.1. Model validation

To determine the validity of the mathematical model, we have used *R*^2^ and p-value of the graphs found from statistical analysis.

For our linear log-log model slope, Y-intercept and *R*^2^ of the data-fitted graphs are calculated.

All the results of our regression analysis on the data can be found in the Tables 1-3. The corresponding graphs can be found in the Extended Data Figures.

**Table 1:**
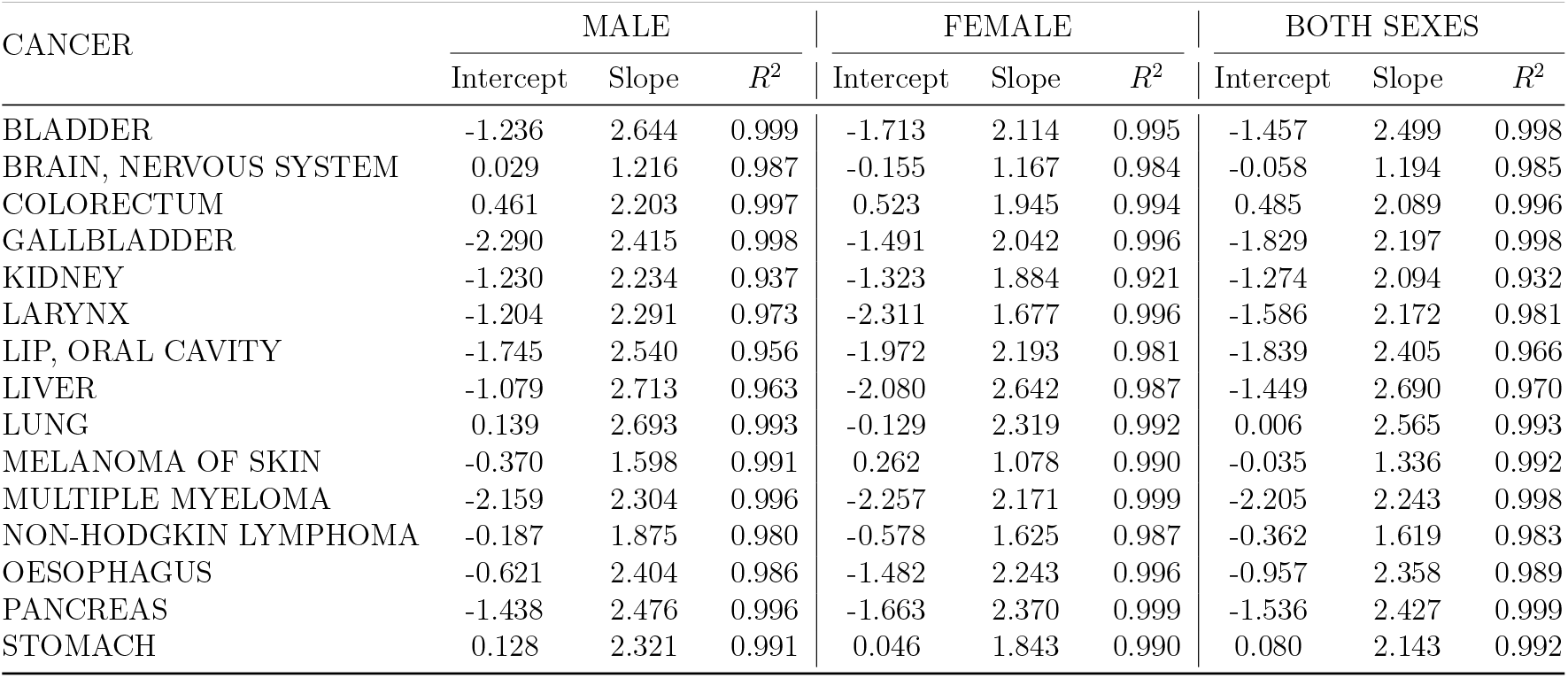
Cancer categories best fitting the linear log-log model.

| CANCER | MALE |  |  | FEMALE |  |  | BOTH SEXES |  |  |
| --- | --- | --- | --- | --- | --- | --- | --- | --- | --- |
| | Intercept | Slope | $R^2$ | Intercept | Slope | $R^2$ | Intercept | Slope | $R^2$ |
| BLADDER | -1.236 | 2.644 | 0.999 | -1.713 | 2.114 | 0.995 | -1.457 | 2.499 | 0.998 |
| BRAIN, NERVOUS SYSTEM | 0.029 | 1.216 | 0.987 | -0.155 | 1.167 | 0.984 | -0.058 | 1.194 | 0.985 |
| COLORECTUM | 0.461 | 2.203 | 0.997 | 0.523 | 1.945 | 0.994 | 0.485 | 2.089 | 0.996 |
| GALLBLADDER | -2.290 | 2.415 | 0.998 | -1.491 | 2.042 | 0.996 | -1.829 | 2.197 | 0.998 |
| KIDNEY | -1.230 | 2.234 | 0.937 | -1.323 | 1.884 | 0.921 | -1.274 | 2.094 | 0.932 |
| LARYNX | -1.204 | 2.291 | 0.973 | -2.311 | 1.677 | 0.996 | -1.586 | 2.172 | 0.981 |
| LIP, ORAL CAVITY | -1.745 | 2.540 | 0.956 | -1.972 | 2.193 | 0.981 | -1.839 | 2.405 | 0.966 |
| LIVER | -1.079 | 2.713 | 0.963 | -2.080 | 2.642 | 0.987 | -1.449 | 2.690 | 0.970 |
| LUNG | 0.139 | 2.693 | 0.993 | -0.129 | 2.319 | 0.992 | 0.006 | 2.565 | 0.993 |
| MELANOMA OF SKIN | -0.370 | 1.598 | 0.991 | 0.262 | 1.078 | 0.990 | -0.035 | 1.336 | 0.992 |
| MULTIPLE MYELOMA | -2.159 | 2.304 | 0.996 | -2.257 | 2.171 | 0.999 | -2.205 | 2.243 | 0.998 |
| NON-HODGKIN LYMPHOMA | -0.187 | 1.875 | 0.980 | -0.578 | 1.625 | 0.987 | -0.362 | 1.619 | 0.983 |
| OESOPHAGUS | -0.621 | 2.404 | 0.986 | -1.482 | 2.243 | 0.996 | -0.957 | 2.358 | 0.989 |
| PANCREAS | -1.438 | 2.476 | 0.996 | -1.663 | 2.370 | 0.999 | -1.536 | 2.427 | 0.999 |
| STOMACH | 0.128 | 2.321 | 0.991 | 0.046 | 1.843 | 0.990 | 0.080 | 2.143 | 0.992 |

**Table 2:** Cancer categories best fitting the convex upwards model.

| SEX | CANCER | $\alpha_0$ | $\alpha_1$ | $\alpha_2$ | $\alpha_3$ | $R^2$ | $k_p$ | $k_q$ | $k_r$ | $k_s$ |
| --- | --- | --- | --- | --- | --- | --- | --- | --- | --- | --- |
| MALE | NASOPHARYNX | -0.54 | 2.797 | 0.73 | 0.14 | 0.9856 | 0.583 | 0.425 | 0.583 | 0.425 |
| MALE | OTHER PHARYNX | -0.485 | 2.844 | 0.47 | 0.33 | 0.9923 | 0.616 | 0.763 | 0.616 | 0.763 |
| FEMALE | NASOPHARYNX | -1.36 | 2.797 | 0.73 | 0.14 | 0.9756 | 0.257 | 0.187 | 0.257 | 0.187 |
| FEMALE | OTHER PHARYNX | -0.642 | 2.5295 | 0.77 | 0.11 | 0.9838 | 0.526 | 1.464 | 0.526 | 1.464 |
| BOTH SEXES | NASOPHARYNX | -0.845 | 2.797 | 0.73 | 0.14 | 0.9847 | 0.429 | 0.313 | 0.429 | 0.313 |
| BOTH SEXES | OTHER PHARYNX | -0.277 | 2.814 | 0.69 | 0.16 | 0.9916 | 0.758 | 0.910 | 0.758 | 0.910 |

**Table 3:** Cancer categories best fitting the concave upwards model.

| SEX | CANCER | $\beta_0$ | $\beta_1$ | $\beta_2$ | $\beta_3$ | $R^2$ | $k_p$ | $k_q$ | $k_r$ | $k_s$ |
| --- | --- | --- | --- | --- | --- | --- | --- | --- | --- | --- |
| MALE | HODGKIN LYMPHOMA | -0.036 | 0.1405 | 0.007 | 2.1 | 0.9956 | 0.964 | 0.007 | 0.964 | 0.007 |
| MALE | LEUKAEMIA | 1.15 | -0.8291 | 0.09 | 2.75 | 0.9941 | 3.158 | 0.284 | 3.158 | 0.284 |
| FEMALE | HODGKIN LYMPHOMA | 0.7843 | -1.6493 | 0.039 | 2.9 | 0.9920 | 2.191 | 0.085 | 2.191 | 0.085 |
| FEMALE | LEUKAEMIA | 0.8742 | -0.8291 | 0.11 | 2.75 | 0.9925 | 2.397 | 0.264 | 2.397 | 0.264 |
| BOTH SEXES | HODGKIN LYMPHOMA | 0.1634 | -0.3739 | 0.025 | 2.1 | 0.9863 | 1.178 | 0.029 | 1.178 | 0.029 |
| BOTH SEXES | LEUKAEMIA | 0.6742 | -0.8291 | 0.11 | 2.75 | 0.9925 | 2.397 | 0.264 | 2.397 | 0.264 |

### 3.2. Extendibility to non-neoplastic conditions

We also found that our model is applicable to any disease process, not only cancers, that satisfy the requirement of progression via discrete rate-limiting steps. Rheumatoid arthritis is modeled as an example of its extendibility. Further tests on data of specific incidences of different chronic diseases are done as potential scope of this research and further validates our mathematical model.

where *α*_0_, *α*_1_, *α*_2_, *α*_3_, *β*_0_, *β*_1_, *β*_2_ and *β*_3_ contain the following terms for the two different assumptions: heterogeneity assumption and age-related effect assumption. *k*_*p*_ and *k*_*q*_ are the respective probabilities of a rate-limiting event to occur with the rate *t*^*p*^ and *t*^*q*^ where p and q are real numbers. *k*_*a*_ is the probability of a rate-limiting event to occur with the rate *t*^*a*^ a is a real number.

## 4. Discussion

### 4.1. Biological interpretation from WHO molecular data

We found that for almost all the cancers fitting the standard linear model, the number of rate-limiting steps ranges from 3-4 on average, except non-Hodgkin lymphoma which requires just one such step under appropriate conditions. This is mostly consistent with the molecular findings described in WHO Cancer “Blue Books” published by the International Agency for Research on Cancer (IARC) as well as different textbooks on cancer pathology and relevant published articles. [11]

For instance, the development of bladder cancer (mainly urothelial carcinoma) requires the driver mutations of genes FGFR3 and RAS along with any one among TP53, Rb or PTEN. [12] Cancers of kidney (mostly renal cell carcinoma) appear to require VHL mutation along with two other genetic alterations that may include EGFR mutation, overexpression of genes associated with cellular adhesion molecules (e.g. E-cadherin) and matrix regulatory proteins (e.g. MMPs, bFGF, VEGF) or rarely p53.

The tumors of the brain and nervous system are quite a heterogeneous group, still confined to the above-mentioned number of driver mutations. For example, most low-grade astrocytomas seem to require mutations in BRAF, RAS and IDH or might follow a different line of progression involving TP53, PDGFR and loss of heterozygosity (LOH) at 22q or p14 promoter methylation. There is a predisposed group of NF1 mutation as well. Whereas high-grade astrocytomas mostly show LOH at 17p, TP53 and PTEN mutations, Glioblastomas on the other hand usually need LOH at 10q, EGFR amplification, p16 deletion and TP53 mutation to manifest. It is to note that only 5% of the glioblastoma progress from low-grade lesions while the rest develop de novo. In case of meningiomas, chromosome 22 deletion consistently disables at least two tumor suppressor genes, one of which is NF2 and the other is yet to be determined. The other significant alteration here is microsatellite instability which is the result of the mutation at least one DNA mismatch repair gene. Genetic studies of several other groups of nervous system tumors like neuronal and mixed neuronal-glial tumors reveal no consistent mutational pattern to date.

Cancers of esophagus show a consistent pattern of mutation in a transcription factor gene (e.g. SOX2), a cell cycle regulator gene (e.g. cyclin D1) and a tumor suppressor gene (commonly TP53) for their pathogenesis. Comparable changes also apply for the cancers of lip, oral cavity, larynx and pharynx. Nasopharyngeal carcinoma will be discussed in a separate context below. A similar repertoire as the esophageal cancers is also noted in the cancers of stomach although consisting of a different set of mutated genes (e.g. CDH1, APC, etc.) along with microsatellite instability. Regarding gastric lymphomas, MLT/BCBL-10 pathway seems to play a pivotal role in molecular pathogenesis 12 which also requires 3-4 driver mutations.

Colorectal cancers are considered the prototypical example to multistep carcinogenesis [2] involving up to ten genes and their mutations as well as microsatellite instability along the line of tumor progression. But Tomasetti et al. (2015) [16] pointed out that only three driver mutations among them at a time are required to produce those carcinomas.

Mutations along the Wnt pathway along with TP53 and cell cycle regulators are necessary for most of the cancers of the liver. Additionally, mutations along the KRAS pathway may be important for cancers of gallbladder and biliary tree. Inactivation of p16 and TP53 along with another rate-limiting step (e.g. BRCA2 mutation) are required for the development of most pancreatic cancers.

Lung cancer has a diverse set of mutations, not necessarily corresponding to its clinical or pathological subtypes. However, the latest WHO classification of lung cancer attempted to formulate a scheme to incorporate the molecular alterations relevant to therapy and prognosis. Across different strata of the scheme, there are 2-3 mutations or rate-limiting steps appear to be necessary to produce the clinical manifestation of the tumor. For instance, mutation of TP53 and Rb genes along with loss of heterozygosity at 3p seems to be a recurring theme in many cases. However, the sufficiency of those alterations is yet to be established as a universal rule.

Mutation of at least one out of five genes (cyclin D1, C-MAF, FGFR3/MMSET, cyclin D3 and MAFB) seems to be a consistent finding in plasma cell myeloma which requires about three such rate limiting steps on average to manifest clinically.

Like lung cancers, a sweeping generalization regarding the rate-limiting steps of skin cancers is also out of reach at the moment. However, a case-by-case analysis reveals similar recurring theme of up to three mutations. Interestingly, non-Hodgkin lymphoma, a highly heterogenous entity, appears to have lower threshold of the required number of mutations, usually up to two rate-limiting steps according to their respective subcategories which is consistent with the model presented in this article.

Some of the cancers, however, do not follow the linear log-log model thus far discussed. To accommodate those entities, we have modeled generalizations assuming either of the two sets of assumptions. We have taken into account the fact that some factors may accelerate or decelerate the rate of progression of the disease with advancing age and named it age-related effect. Other assumption is based on the variation of the required number of rate-limiting steps amongst affected population due to heterogeneity of the condition under consideration. Four such entities which include cancers of nasopharynx, thyroid, leukemia and Hodgkin lymphoma, are modeled under the aforementioned generalization.

Nasopharyngeal carcinoma shows up to five mutations including DN-P63, P27, cyclin D1 and BCL-2 associated with its development and manifestation which is consistent with our convexity-upwards log-log model. We also predict age-related deceleration of its carcinogenesis due to some yet undiscovered factor, possibly associated with Epstein-Barr virus (EBV) infection. Thyroid neoplasms are often associated with the mutations of RET/PTC, RAS, BRAF and PTEN which is quantitatively consistent with our model. The heterogeneity of thyroid neoplasms may account for the necessity of heterogeneity assumptions used in its generalization. Interestingly, the number of mutational steps predicted for leukemia and Hodgkin lymphoma is comparable with that of non-Hodgkin lymphoma, except for the requirement of additional generalizing assumptions required for the former pair. All three categories of those hematolymphoid neoplasms require relatively lower number of rate-limiting steps. However, heterogeneity and/or age-related acceleration of carcinogenesis may play more decisive role in leukemia and Hodgkin lymphoma according to our model.

### 4.2. Significance of different quantities of the Standard Model

#### 4.2.1. Effect of changing the scale of t

Since *k*_*n*_ denotes the probability per unit of time, the scale for time affects its value. For instance, if the *t* is scaled to *w* times, i.e., two adjacent time intervals differ by *w* units, then every *k*_*n*_ will need to be replaced by 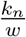 and the right-hand side of the

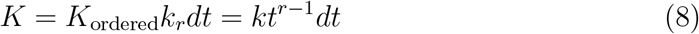

would have an extra ‘constant’ term *w*^*r−*1^ at its denominator. Subsequent steps of the calculation would show that this change would only affect the term ln *k* which is the intercept of the final log-log linear form,

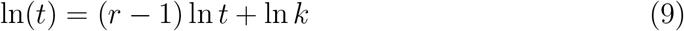

but does not affect the slope (*r* − 1). So, for the sake of simplicity, we would use unity as the log scale for time. Hence, age classes 1, 2, 3 are used in all of our graphs. If any cancer has zero incidence at the initial age class (0-14 years) for both male and female then the subsequent age class is designated as class 1 and consecutively onwards. This is for avoiding structural zeroes in log-log plot since logarithm of zero is undefined.

#### 4.2.2. Effect of male-to-female ratio of age-specific incidence rate on slope

For reasons explained later, almost all of the cancers show two interesting features: firstly, the three linear plots (male, female and both sexes) for each cancer have somewhat unequal slopes, and secondly, the linear plot for both sexes of a given cancer shows a slope which has a value that is within the range of the slopes obtained from the separate linear plots for male and female population of the same cancer. For example, in the case of cancers of brain and nervous system, the best-fit (*R*^2^ = 0.98) linear plot gives 1.16 and 1.21 as slopes for females and males, respectively, while the plot for both sexes has a slope of 1.19.

Let us explore the second feature first. It is tempting to assume that this happens simply because of the fact that, for a given age class of a cancer, the age specific incidence rate of both sex (*I*_m_) group is by definition equal to the arithmetic mean of the rates of the male (*I*_m_) and the female (*I*_f_) groups,

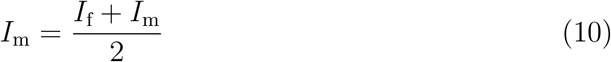

But the actual reason is a bit more non-trivial.

**Slope of the female line**,

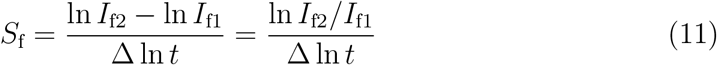

where, (ln *t*_1_, ln *I*_f1_) and (ln *t*_2_, ln *I*_f2_) are two points on the line and Δ ln *t* = ln *t*_2_ − ln *t*_1_ *>* 0.

Similarly, **Slope of the male line** over the same abscissae,

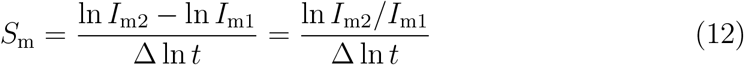

And the line for both sexes likewise has the slope,

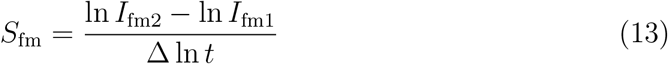

The above equation can be rewritten as,

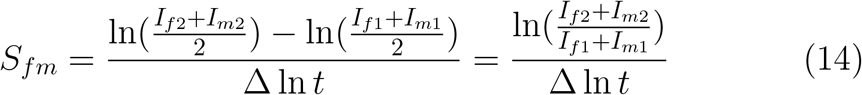

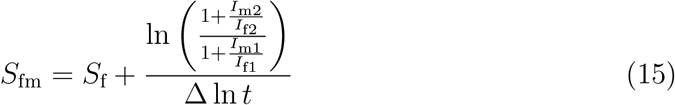

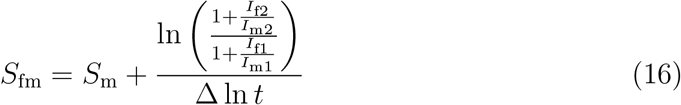

Let, *I*_f_ *< I*_m_. Then the difference between *I*_m_ and *I*_f_ would be greater for higher values of *t* than its lower values. Therefore, 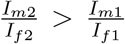 and thus, the second term on right hand side of Equation 15 must be positive, ensuring *S*_fm_ *> S*_f_. By similar deduction from Equation 16, *S*_fm_ *< S*_m_ is also guaranteed. So, *I*_f_ *< I*_m_ implies *S*_m_ *> S*_fm_ *> S*_f_. Conversely, *I*_f_ *> I*_m_ implies *S*_m_ *< S*_fm_ *< S*_f_. Both the scenarios are consistent with the aforementioned second feature of the plots. And if *I*_f_ = *I*_m_ then the second terms of both Equation 15 and Equation 16 become zero and thus, *S*_m_ = *S*_m_ = *S*_f_, which is not observed in any of our plots due to the persistent inequality of the incidence rates between sexes at all instances; without exception.

#### 4.2.3. Effect of the rate of limiting events on slope and intercept

In the previous section we explored why the slope of the both-sexes plot happens to be intermediate between the slopes of the plots for male and female drawn separately. Now we will look into the possible explanations for the inequality of the slopes for male and female in the first place. Beyond the trivial reasons like experimental error, there might be fundamental biological attributes responsible for the observed difference. One of the possibilities is the violation of assumption 7 where rate of any or few or all of the rate-limiting events is a function of time. If the rate of the *n*-th mutation per unit of time (*k*_*n*_) is exponentially proportionate to time at its *h*-th power (*t*^*h*^) then

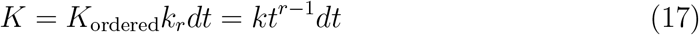

will have an extra term *t*^*h*^ multiplied to it (*h* ∈ ℝ), eventually rendering

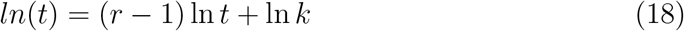

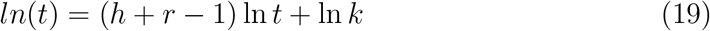

which will be indistinguishable from the Standard Model when the coefficients are numerical as in a real plot, but must be interpreted differently because the slope (*h*+*r* − 1) no longer denotes the number of rate-limiting steps (*t*) alone. The problem is, just from the linear plot-fitting exercise alone, one cannot be certain if the slope equals to (*h*+*r* − 1) or just (*r* − 1). But from the molecular data on tumor progression, two inferences can be incorporated as extended assumptions when assumption 7 does not hold:

8. Sex-associated differences in the slopes for a given cancer may be attributed to different values of the exponent of the aforementioned exponential mutation rate between sexes and not to the possibility of having different numbers of rate-limiting events for the same cancer in males and females.
9. Assumption 8 may be repurposed and incorporated to quantify geographical and/or ethnic as well as environmental and lifestyle-related differences of cancer progression.

## 5. Conclusion

By uniting mathematical and biological concepts, this framework has the potential to transform how cancer and other diseases are studied. Its contributions can extend to the development of more effective diagnostic methods and therapeutic strategies, ultimately advancing both theoretical understanding and practical applications in oncology. This work represents a significant step toward a more integrated, data-driven approach to tackling complex diseases.

---

**Figure 1.**
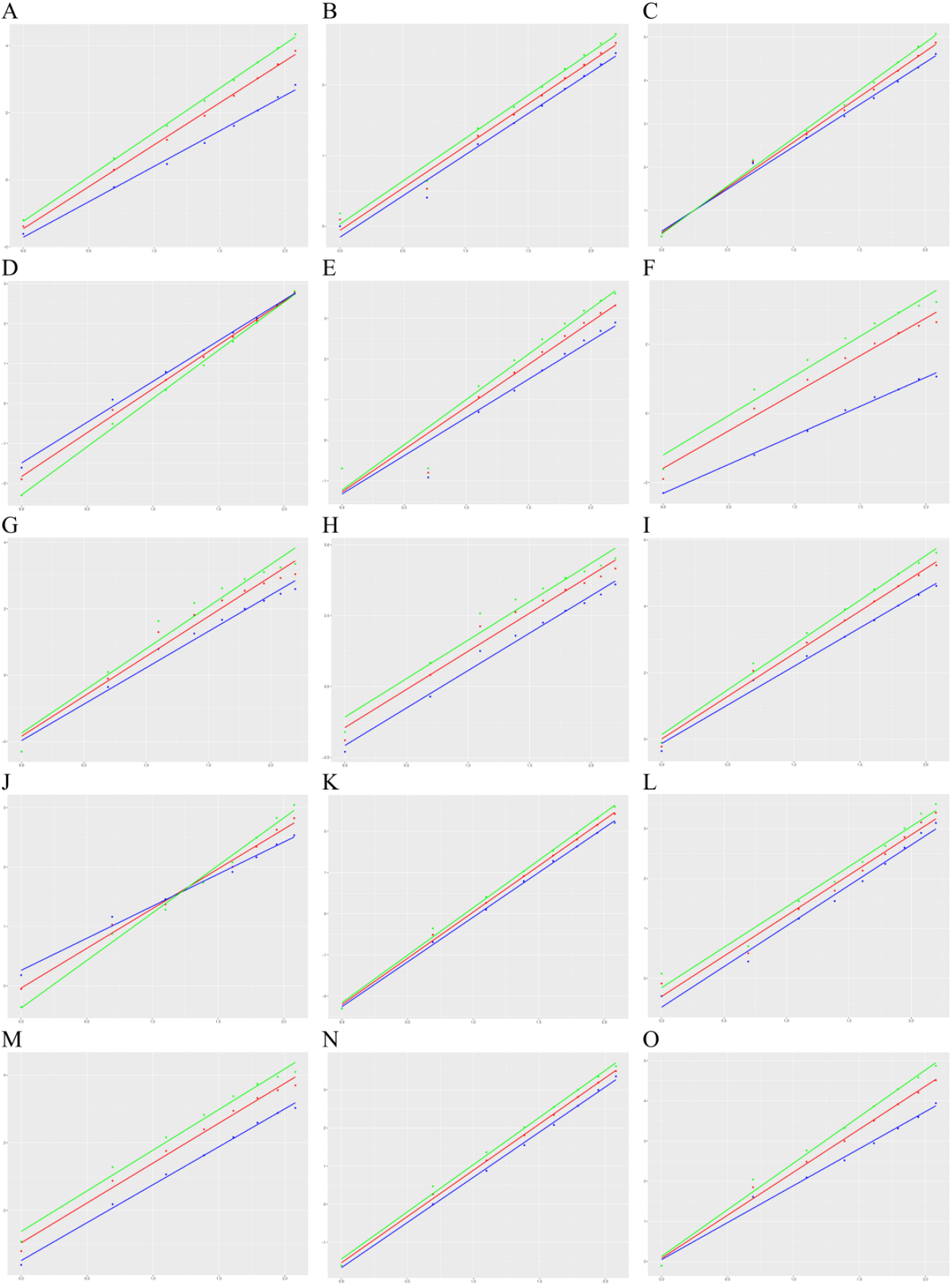
Cancer categories best fitting the linear log-log model. The horizontal and vertical axes show age class and incidence, respectively, both in log scale. Blue, green, and red lines and dots represent the regression lines and data points for female, male, and both sexes, respectively. A, bladder cancer; B, cancer of brain and nervous system; C, colorectal cancer; D, gallbladder cancer; E, kidney cancer; F, cancer of larynx; G, cancer of lip and oral cavity; H, liver cancer; I, lung cancer; J, skin melanoma; K, multiple myeloma; L, non-Hodgkin lymphoma; M, cancer of esophagus; N, pancreatic cancer; O, stomach cancer.

**Figure 2.**
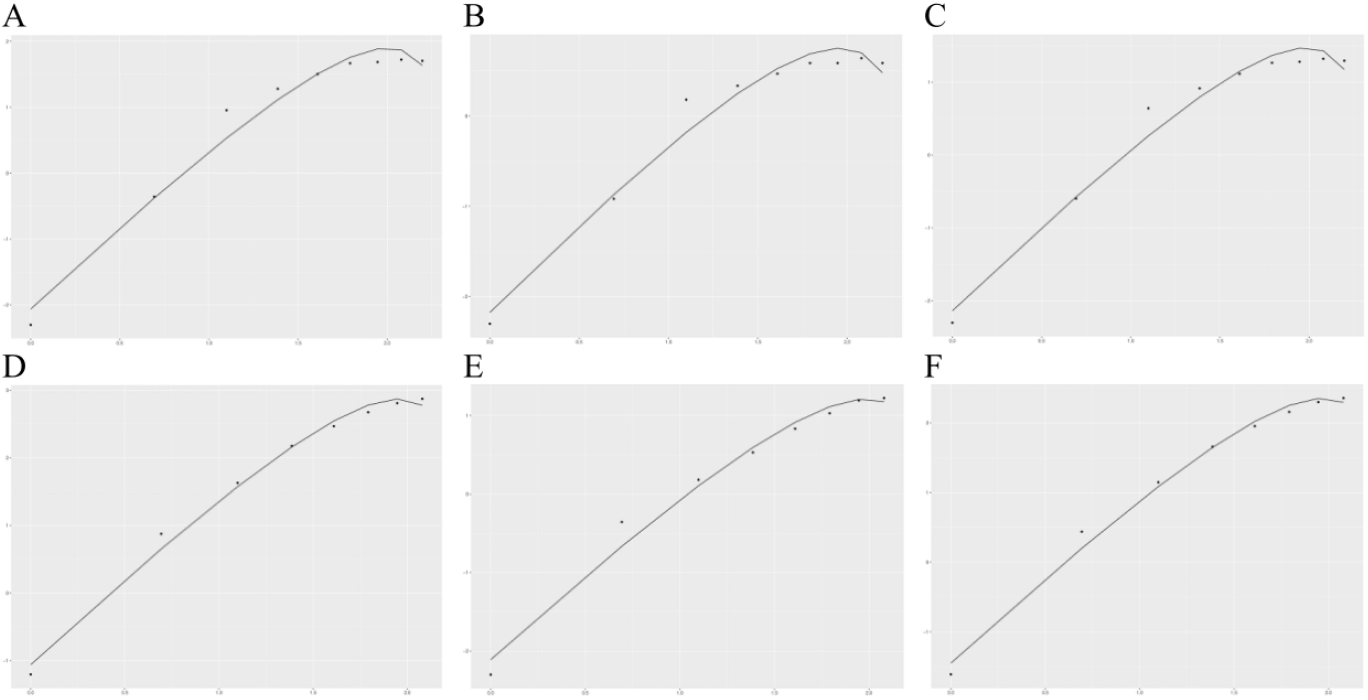
Cancer categories best fitting the convex upwards model. The horizontal and vertical axes show age class and incidence, respectively, both in log scale. A, cancer of nasopharynx in male; B, cancer of nasopharynx in female; C, cancer of nasopharynx in both sexes; D, cancer of other parts of the pharynx in male; E, cancer of other parts of the pharynx in female; F, cancer of other parts of the pharynx in both sexes.

**Figure 3.**
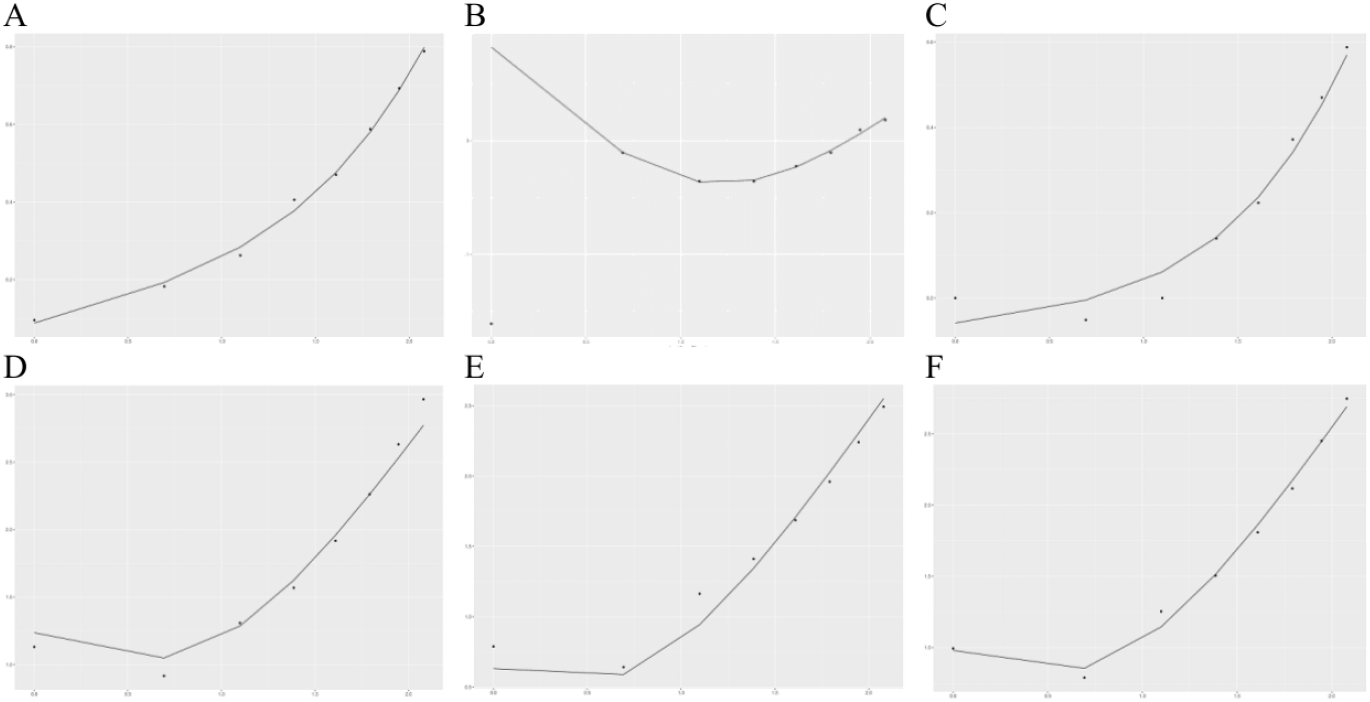
Cancer categories best fitting the concave upwards model. The horizontal and vertical axes show age class and incidence, respectively, both in log scale. A, Hodgkin lymphoma in male; B, Hodgkin lymphoma in female; C, Hodgkin lymphoma in both sexes; D, leukemias in male; E, leukemias in female; F, leukemias in both sexes.

**Figure 4.**
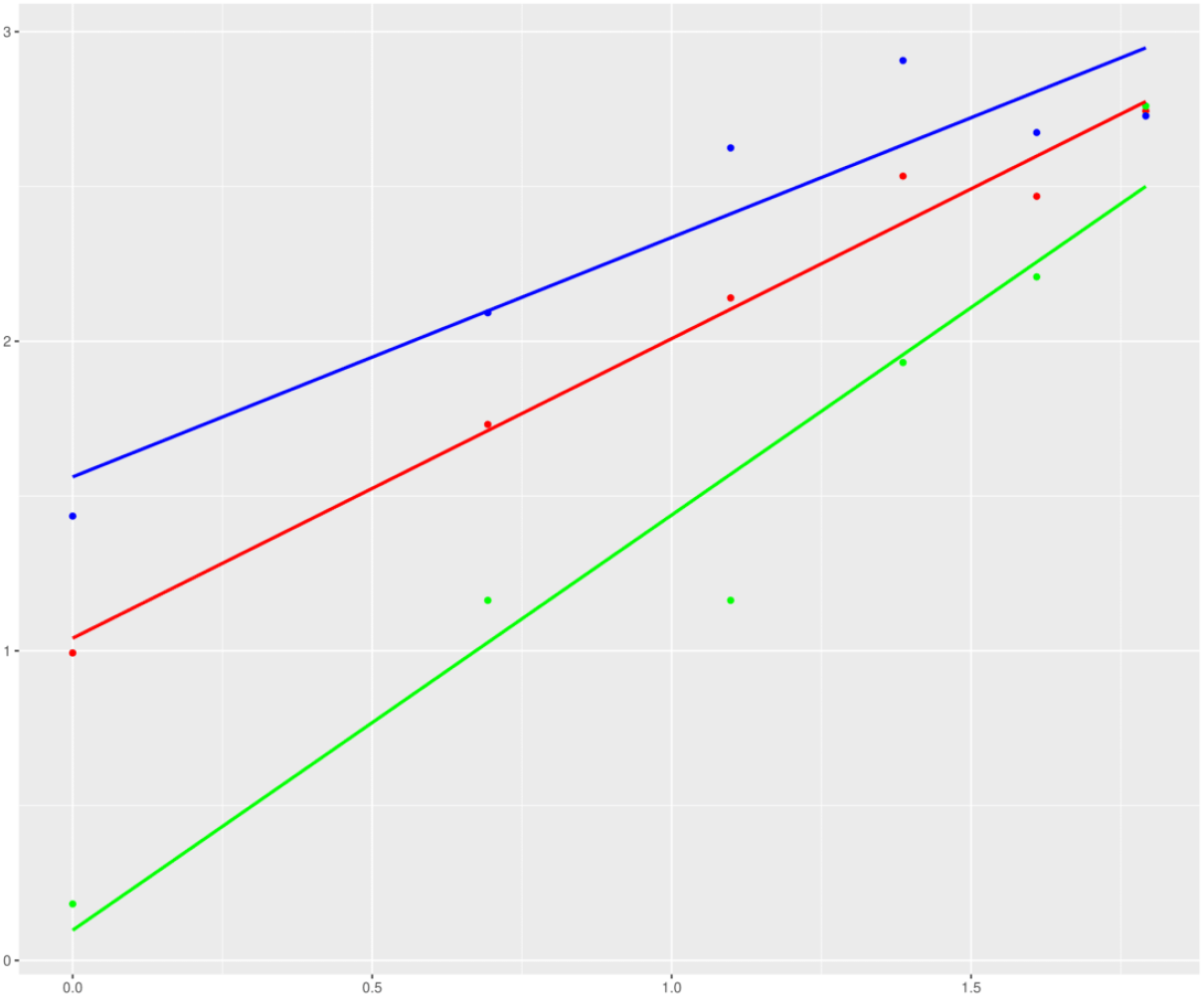
The linear log-log model has been extended to predict genetic and pathophysiological rate limiting events in rheumatoid arthritis, a non-neoplastic condition. The horizontal and vertical axes show age class and incidence, respectively, both in log scale. Blue, green, and red lines and dots represent the regression lines and data points for female, male, and both sexes, respectively. Age groups of 21-30, 31-40, 41-50, 51-60, 61-70, and >70 years have been denoted by the age classes 1-6, respectively. The equations of the regression line for male: log (incidence)= 0.0975 + 1.3411 × log (age class), R^2^ = 0.9377; for female: log (incidence)= 1.5619 + 0.7735 × log (age class), R^2^ = 0.8668; for both sexes: log (incidence)= 1.0407 + 0.9677 × log (age class), R^2^ = 0.9787.

