## Appendix A for "Mathematical bridge between epidemiological and molecular data on cancer and beyond"

### Appendix A. Derivations

#### *Appendix A.1. Derivation of the standard (linear) model*

Standard model of multistep carcinogenesis refers to the mathematical as well as statistical framework which allows us to simulate, test and predict the number of rate-limiting steps for a given cancer in order to gain mechanistic insight into carcinogenesis through communication between epidemiological and molecular data. We will derive the standard model in the following steps.

1. Derivation of the probability of a given number of sequential rate-limiting events to occur within a given duration of time.
2. Derivation of the probability of $r$-th rate-limiting events to occur in an infinitesimal time interval after $r-1$ such events have already occurred.
3. Evaluation of the function that takes time as input and gives probability as output. Here, the probability means the chance that an individual of the age group denoted by the time does not manifest a cancer yet, all other quantities being constant.
4. Derivation of the formula for age-specific incidence rate of cancer.

**Step 1**

If $r-1$ rate-limiting events occur with $k_1, k_2, \ldots, k_{r-1}$ probability per unit of time then the probability of each of those occurring in course of time $t$ would be $k_1 t, k_2 t, \ldots, k_{r-1} t$, respectively, provided that the values of $k_1, k_2, \ldots, k_{r-1}$ are small enough to allow the probabilities $k_1 t, k_2 t, \ldots, k_{r-1} t$ to scale for a value of $t$ large enough to accommodate typical average human lifespan (Assumption 2). The probability of $r-1$ rate-limiting events to occur in any order during a period of $t$ would thus be,

$$K_{\text{unordered}} = (k_1 t)(k_2 t)\ldots(k_{r-1} t) = (k_1 k_2 \ldots k_{r-1}) t^{r-1} \quad \text{... Equation 1}$$

Since the sequence of mutation has to be ordered, we are interested in the probability of only one out of $(r-1)!$ possible arrangements of rate-limiting event (Assumption 3). Therefore the probability of $r-1$ mutations to occur in a given order during a period of $t$ would be,

$$K_{\text{ordered}} = \frac{K_{\text{unordered}}}{(r-1)!} = \frac{(k_1 k_2 \ldots k_{r-1}) t^{r-1}}{(r-1)!} \quad \text{... Equation 2}$$

**Step 2**

If the $r$-th step, the final rate-limiting event of the impending doom occurs with a probability $k$, per unit of time within a short interval $(t, t + dt)$ then the actual probability of that event would be $k \cdot dt$.

Let the aggregated ordered rate of occurrence of events per unit of time or the probability of all $r$ events to occur in a specific order within the given unit of time,

$$k = \frac{k_1 k_2 \ldots k_{r-1} k_r}{(r-1)!} \quad \text{... Equation 3}$$

The total probability of $(r - 1)$ rate-limiting events to occur in a given order during $t$ followed by the occurrence of $r$-th event during $dt$ would thus be,

$$K = K_{\text{ordered}} k_r dt = \left( \frac{k_1 k_2 \ldots k_{r-1} \cdot t^{r-1}}{(r-1)!} \right) k_r \cdot dt = k \cdot t^{r-1} \cdot dt \quad \text{... Equation 4}$$

**Step 3**

Let $P(t)$ be the probability that an individual of age $t$ has not manifested a given type of cancer yet. Then the total probability of randomly choosing such an individual who eventually develops cancer after a short interval of $dt$ would be,

$$KP(t) = P(t)k \cdot t^{r-1} \cdot dt \quad \text{... Equation 5}$$

This quantity is equivalent to the frequency of individuals of age $t$ developing cancer after they have reached that age and before they reach the age of $(t + 1)$, which is by definition, age specific incidence rate at age $t$. Are we done then? Unfortunately, no! For the sake of mathematical rigor, we have to get rid of the $dt$ term. Besides, in order to obtain a model-worthy form of the formula for age specific incidence rate we have to play a few more mathematical tricks.

Now, let us take random individuals from all possible age groups from 0 to $t$ and calculate their total probability of developing cancer after a short while of $dt$,

$$\sum_{0}^{t} KP(t) \approx \int_0^t P(t) k t^{r-1} \, dt \quad \text{... Equation 6}$$

Please note that their individual probabilities are disjoint events. The integral can also be thought of as a cumulative frequency of individuals between ages 0 and $t$ who developed cancer while remaining at their respective age groups. If a population begins with a frequency (or probability) of $P(0)$ which denotes the individuals at age 0 without the manifestations of cancer and then proceeds up to age $t$ while 'losing' individuals who succumb to cancer at each successive age groups then,

$$P(t) = P(0) - \int_0^t P(t) k\, t^{r-1}\, dt \quad \text{... Equation 7}$$

Since $P(0)$ is a constant term by virtue of being the $0^{th}$ term of a function, differentiating both sides with respect to $dt$ gives us,

$$\frac{d}{dt}P(t) = 0 - P(t)k\, t^{r-1} \Rightarrow \frac{dP(t)}{P(t)} = -k\, t^{r-1}\, dt$$

Integrating both sides,

$$\ln P(t) = -\frac{k\, t^r}{r} + C_0 \Rightarrow P(t) = e^{-\frac{k\, t^r}{r + C_0}} = C_1 e^{-\frac{k\, t^r}{r}}$$

[where $C_0, C_1$ are constants; let $C_1 = e^{C_0}$]

Since at $t = 0$, $P(0) = C_1$,

$$P(t) = P(0) e^{-\frac{k\, t^r}{r}} \quad \text{... Equation 8}$$

**Step 4**

$P(t)$ is a cumulative distribution function (CDF) that expresses the total probability of not developing cancer in the age range of 0 to $t$. It can be thought of as a survival function in a sense that, it denotes the portion of the population at age $t$ that has 'survived' by not developing cancer yet. More rigorously, if we take a CDF, $F(t)$ to indicate the probability of developing cancer in the age range of 0 to $t$ then by definition of survival function,

$$P(t) = 1 - F(t) \quad \text{... Equation 9}$$

We want to determine age specific incidence rate, the probability of developing cancer at any given short interval $(t, t + dt)$ which is nothing but the hazard function of $F(t)$, since hazard function is defined as the probability of an event to occur in a short interval. Since the probability density function (PDF) of $F(t)$ is given by the derivative of $F(t)$ with respect to $dt$ and hazard function is the ratio of PDF to survival function, age specific incidence rate at age $t$ would be,

$$I(t) = \frac{\frac{d}{dt}F(t)}{1 - F(t)} = \frac{\frac{d}{dt}(1 - P(t))}{P(t)} = \frac{P(0)kt^{r-1}e^{-kt^r/r}}{P(0)e^{-kt^r/r}} = kt^{r-1} \quad \text{... Equation 10}$$

This nice and simple equation can be reframed using logarithm as,

$$\ln I(t) = (r-1)\ln t + \ln k \quad \text{... Equation 11}$$

*Appendix A.2. Derivation of the non-linear models from heterogeneity assumption*

If we replace $k = k_p t^p + k_q t^q$ in Equation 4 with the following assumption, $k_p$ and $k_q$ are the respective probabilities of a rate-limiting event to occur with the rate $t^p$ and $t^q$ where $p$ and $q$ are real numbers, then the equation becomes,

$$K = (k_p t^p + k_q t^q)t^{r-1}dt = k_p t^{r+p-1}dt + k_q t^{r+q-1}dt \quad \text{... Equation 12}$$

The rationale behind this assumption is the diversity of factors and mechanisms underlying the heterogeneous groups of disorders clumped under a single label, e.g. leukemia. The simplest form that could accommodate such a situation is a two-factor model [? ] [? ] which is incorporated in the assumption above. If we follow through the calculations according to the derivation of the standard model, then this new assumption ultimately leads to,

$$I(t) = k_p t^{r+p-1} + k_q t^{r+q-1} = k_p t^{r+p-1}\left(1 + \frac{k_q}{k_p}t^{q-p}\right) \quad \text{... Equation 13}$$

Taking log on each side,

$$\ln I(t) = \ln k_p + (r+p-1)\ln t + \ln\left(1 + \frac{k_q}{k_p}t^{q-p}\right) \quad \text{... Equation 14}$$

This is the general form of the equation for log-log plots with upwards concavity. If instead, the initial two-factor term (Equation 12) was assumed to be $k = k_p t^p - k_q t^q$ then the ultimate log-log form would become,

$$\ln I(t) = \ln k_p + (r+p-1)\ln t + \ln\left(1 - \frac{k_q}{k_p}t^{q-p}\right) \quad \text{... Equation 15}$$

This is the general form of the equation for log-log plots with upwards convexity. An additional assumption of $k_q \leq k_p$ would allow for $0 \leq \frac{k_q}{k_p} \leq 1$ to be true while validating the models using real world data. This would help normalize the dataset without compromising the rigor of the method. Also, putting $k_q = 0$ regresses the equations back to the standard linear form.

*Appendix A.3. Derivation of the non-linear models from age-related effect assumption*

If we take into consideration that some factors accelerate or decelerate the rate of tumor progression with increasing age then we could formulate an alternate set of assumptions for non-linear models.

If we add a factor $k_a t^a$ to account for the age-related acceleration in Equation 4 with the following assumption, $k_a$ is the probability of a rate-limiting event to occur with the rate $t^a$ a is a real number, then the equation becomes,

$$K = kt^{r-1}dt + k_a t^a dt \quad \text{... Equation 16}$$

If we follow through the calculations according to the derivation of the standard model, then this new assumption ultimately leads to,

$$I(t) = kt^{r-1} + k_a t^a = kt^{r-1}\left(1 + \frac{k_a}{k} t^{a-r+1}\right) \quad \text{... Equation 17}$$

Taking log on each side,

$$\ln I(t) = \ln k + (r-1)\ln t + \ln\left(1 + \frac{k_a}{k} t^{a-r+1}\right) \quad \text{... Equation 18}$$

This is the general form of the equation for log-log plots with upwards concavity. Similarly, subtracting an age-related deceleration factor $k_d t^d$ from Equation 4 would ultimately result in,

$$\ln I(t) = \ln k + (r-1)\ln t + \ln\left(1 - \frac{k_d}{k} t^{d-r+1}\right) \quad \text{... Equation 19}$$

This is the general form of the equation for log-log plots with upwards convexity. Since the terms $k$, $k_a$, and $k_d$ are probabilities, the fractional terms inside the third log term of Equation 18 and Equation 19 would always be within 0 and 1, inclusive. Also, if there is no age-related acceleration ($k_a = 0$) or deceleration ($k_d = 0$) or if they are small enough to be ignored then the non-linear form regresses to the standard linear form.
